# Automated generation of a gene perturbation transcriptomic atlas using large language models

**DOI:** 10.64898/2026.08.08.743502

**Authors:** Jamie Soul, David A Young

## Abstract

Public transcriptomic repositories contain thousands of gene perturbation experiments, a valuable resource for understanding gene function, but perturbation metadata are not structured, which blocks systematic reuse. Existing perturbation atlases depend on expert manual curation, so they are costly to maintain and infrequently updated, while automated grouping approaches neither identify which samples form the perturbation arm nor recover the perturbed gene.

Here we develop an automated pipeline that uses large language models to find single-gene perturbation experiments in NCBI-GEO and reconstruct their case-control sample groupings, along with the perturbed gene, perturbation type and cell line as structured, ontology-normalised fields. We manually curated 3,300 GEO experiments with sample-level case-control assignments and release these as an open benchmark (2,400 training, 600 validation, 300 temporally held-out test). Reasoning models and task-specific finetuning substantially improved identification of valid perturbation groups, with the best model reaching precision 0.925 and recall 0.836 on the test set. Applied at scale, the pipeline generated an atlas of 6,802 gene perturbation expression signatures from 4,453 GEO experiments, covering 2,907 uniquely perturbed genes.

An R package, perturbMatch, supports exploration of the atlas and querying of user-supplied expression signatures against it using similarity scoring, so users can identify experiments that recapitulate a transcriptional state of interest.

## Introduction

Gene perturbation experiments such as overexpression, knockdown and knockout are widely used to characterise the downstream signalling effects of a target gene (Dixit et al. 2016). For instance, knockdown of a potential drug target in disease can be used to generate gene expression signatures that identify candidate disease mechanisms and inform therapeutic hypotheses (Barbosa et al. 2024). Thousands of such experiments have been deposited in public repositories such as NCBI-GEO using microarray and RNA-Seq platforms representing a large and growing resource for understanding gene function (Clough et al. 2023).

Re-analysis of these experiments allows exploration of differentially expressed genes following perturbation, enables matching of expression signatures to generate hypotheses for functional follow-up, and supports meta-scientific questions about which genes have been studied and which remain largely uncharacterised (Richardson et al. 2024; L. Zhang et al. 2020; Dudley et al. 2011). Such analyses can reveal research gaps and nominate novel avenues of investigation (Stoeger et al. 2018).

Despite the potential of this resource, large-scale analysis is impeded by the quality and availability of metadata (Gonçalves and Musen 2019; Huang et al. 2025). Depositors to NCBI-GEO are not required to annotate case and control samples, and the resulting metadata are unstructured and non-uniform. Importantly, there is no designated field for the experimental variable, and control samples may be labelled in any number of ways with no standardisation enforced e.g. ctrl, control, wild-type, wt (Wang et al. 2018; Bernstein et al. 2017). For more complex experimental designs there may be multiple plausible control groups, for example non-transfected cells versus an empty vector control, and perturbations performed across several cell lines. Resources such as Recount2, ARCHS4 and more recently GEO itself have made raw data available as count-based matrices suitable for downstream differential expression analysis, but these do not provide the case-control sample groupings or perturbed gene identity essential to compute gene expression signatures and use for downstream tasks (Collado-Torres et al. 2017; Lachmann et al. 2018).

Existing gene perturbation expression atlases such as CREEDS, the Gene Perturbation Atlas, GPSAdb and PerturbAtlas have been assembled through expert manual curation (Wang et al. 2016; Xiao et al. 2015; Y. Zhang et al. 2025; Guo et al. 2026). While manual curation produces accurate annotations, it is resource intensive and these perturbation databases may not be regularly updated. Earlier computational approaches have attempted to address this challenge. GEOracle applies natural language processing and supervised machine learning to GEO free-text metadata to classify perturbation experiments and match treatment and control samples, presented via an interactive R/Shiny interface that requires user verification of each experiment (Djordjevic et al. 2019). More recently, RummaGEO uses K-means clustering of embedded sample metadata fields to automatically group GEO samples for differential expression analysis (Marino et al. 2024). While scalable, this approach does not directly identify which group represents the perturbation versus the control, nor does it extract the perturbed gene identity or perturbation type required to build a gene perturbation atlas. Furthermore, clustering-based approaches may not handle the range of challenges present in GEO deposits, including inconsistent perturbation terminology and experimental designs where the correct grouping depends on interpreting unstructured experimental context. Large language models (LLMs) offer a more flexible approach to this problem and have been successfully applied to automate complex curation tasks requiring domain knowledge, owing to their ability to process unstructured text (Zhao et al. 2026; Shintani et al. 2026). LLMs have recently been used to annotate Sequence Read Archive (SRA) entries with sample-level metadata including cell line, but this has not been extended to identifying the case-control groupings required to compute gene expression signatures from perturbation experiments (Cinquin 2024; Ikeda et al. 2025). Unlike sample-level entity extraction, case-control grouping requires inferring the intended contrast across a whole series and choosing between plausible control arms.

Here we address this gap with four contributions. First, we construct and openly release a benchmark of 3,300 manually curated GEO experiments annotated with sample-level case-control groupings, perturbed gene, perturbation type and cell line, including negative examples and a temporally held-out test set. To our knowledge this is the first benchmark for perturbation case-control reconstruction and it provides a means of evaluating future approaches on this task. Second, we use it to benchmark open-weight LLMs spanning instruction-tuned, reasoning and finetuned variants, showing that models of ≤30B parameters running on a single GPU reach precision approaching 0.93, removing dependence on commercial APIs. Third, we couple LLM extraction with HGNC and EFO normalisation to yield structured, ontology-mapped metadata fields that are not systematically provided by existing automated approaches. Fourth, we apply the pipeline across GEO to generate an atlas of gene perturbation signatures and provide an R package supporting signature querying, enabling researchers to identify perturbations that recapitulate or reverse a transcriptional state of interest. Implemented in Nextflow, the pipeline is designed to be re-run so the resource can be updated with new GEO submissions.

## Methods

### Identification of candidate gene perturbation experiments from NCBI-GEO

Human RNA-Seq and microarray gene expression series in GEO were identified using a set of search terms (conducted 29^th^ August 2025) describing genetic perturbations: RNA interference (siRNA, shRNA, RNAi), genome editing or knockout approaches (CRISPR, KO, knockout), antisense oligonucleotides (ASO), and gene overexpression or depletion. Experiments were filtered to retain those with a minimum of 4 samples (to conduct differential expression analysis) and maximum of 40 samples (deemed unlikely to be gene perturbation studies).

### Data curation and preparation

Gene perturbation experiments from the Gene Expression Omnibus (GEO), including both microarray and RNA-Seq platforms, were manually curated to build a benchmarking dataset. Groups consisting of perturbation samples (e.g. CRISPR knockout, siRNA, or overexpression) and their corresponding control samples generated under the same experimental conditions were identified. For each group, we recorded the GEO accession, cell line, perturbed gene, and perturbation type. To accelerate curation, annotations from existing manually curated resources including CREEDS and the Gene Perturbation Atlas were incorporated where available, with all entries reviewed and filtered to ensure consistency with our curation criteria.

Groups were intended to capture the direct transcriptional effects of single-gene perturbations and were therefore restricted to groups with perturbation of a wild-type single gene performed under baseline culture conditions (e.g. normoxia and standard growth media). Experiments involving differentiation, stress induction, or additional treatments and small-molecule perturbations were excluded.

Experiments with no identifiable perturbation groups were randomly sampled from GEO and included as negative examples to evaluate false positive rates. Datasets released after 2023 were reserved as an independent test set to assess model generalisation and potential distributional shift over time. The remaining pre-2023 data were split 80/20 into training and validation sets for model development and hyperparameter tuning.

### Selection of LLMs

Open-weight large language models were selected from Hugging Face to represent diverse model families and parameter scales, including both instruction-tuned and reasoning-optimised variants (Wolf et al. 2020). Models were restricted to a maximum of 30 billion parameters (30B) to maintain computational feasibility for large-scale inference on a single NVIDIA L40S GPU.

### LLM prompting

LLM prompts were designed to identify case-control groups from GEO gene expression series according to the curation criteria described above (Supplementary Note 1). The system prompt specified the task, required output format (YAML), and guidelines for extracting key metadata fields including cell line, perturbation method, and target gene. Valid groups were required to comprise human samples from a single consistent cell line, a single perturbation method targeting one gene, and a minimum of 2 case and 2 control samples. Control samples were assigned by prioritising negative or mock perturbation conditions. Non-transcriptomic data and multi-gene perturbations were excluded. The user prompt included two annotated input-output examples to illustrate the expected output structure and application of the curation criteria.

### LLM finetuning

Finetuning was performed using the Unsloth framework with LoRA adapters applied to instruction-tuned Qwen models (4B and 30B parameters) (Hu et al. 2021). Models were loaded in 4-bit quantisation with gradient checkpointing to reduce GPU memory requirements. The training dataset consisted of curated instruction-response pairs covering both valid perturbation groups and negative examples with no identifiable group. Data were tokenised and formatted using the SFTTrainer framework. Models were optimised using the AdamW 8-bit optimiser with linear learning rate scheduling, weight decay regularisation, and a dynamic warmup period. A hyperparameter search over learning rate, LoRA rank, and random seed was conducted, with the final model selected based on the highest mean F1 score across three random seeds on the validation set.

### Model inference and response parsing

Model inference was performed using the Unsloth framework. Models exceeding 27B parameters were loaded with 4-bit quantisation. Model-specific chat templates were applied prior to generation and outputs were generated using temperature = 0, with a maximum of 12,000 new tokens per sample to accommodate the reasoning traces produced by reasoning models. Chain-of-thought reasoning traces were removed from responses prior to downstream parsing and evaluation.

### Gene normalisation

Gene symbols were normalised to official HGNC-approved symbols using a reference database of human genes constructed from the NCBI Gene Entrez API, containing official symbols, gene descriptions, and synonyms. Predicted gene symbols were initially matched by exact lookup against this database. For symbols without an exact match, fuzzy matching was performed against known synonyms using RapidFuzz (token set ratio; score cutoff ≥70) returning up to five candidates (Bachmann et al. 2025). Candidates were passed to Magistral-Small alongside the GEO experiment metadata to select the most appropriate official symbol. Where no suitable candidates were identified, the original predicted symbol was retained.

### Cell line normalisation

Cell line annotations were normalised to the Experimental Factor Ontology (EFO) using text2term (v4.6.0), matching against the cell-relevant branches of EFO supplemented with Cellosaurus for lines not represented in EFO (Gonçalves et al. 2024; Bairoch 2018; Malone et al. 2010). Annotations were first matched by exact synonym lookup and otherwise by fuzzy matching, with high-confidence matches (score ≥ 0.8) accepted automatically. Ambiguous terms were passed to Magistral-Small alongside the GEO experiment title and summary to select the best candidate or reject all.

### Evaluation of performance

Predicted case and control sample groups were compared against manually curated ground truth annotations. For each prediction, Jaccard similarity was computed between predicted and true GSM identifiers for case and control samples separately, and the mean taken as an overall overlap score. Each prediction was matched to the ground truth group maximising this score, and a prediction was considered correct only when both sets exactly matched (overlap score = 1).

Precision was defined as the proportion of predicted groups that were correct, recall as the proportion of true groups recovered, and F1 as their harmonic mean. True negatives were defined as experiments where no groups were predicted and none were present in the ground truth. Predicted groups absent from the ground truth were treated as false positives, and unrecovered ground truth groups as false negatives.

For correct group predictions, additional metadata fields were evaluated independently: perturbed gene symbol, cell line, and perturbation type. All metrics were computed per model across the validation and held-out test sets. For final annotation, both finetuned Qwen3-30B-A3B-Instruct and Magistral-Small were applied to the full dataset. Case-control groupings identified by either model were retained. Where the two models agreed on grouping but disagreed on its associated metadata Magistral-Small annotations were used.

### RNA-Seq processing

Raw count matrices were downloaded from NCBI-GEO for annotated RNA-Seq accessions where available. Samples were assigned to case and control groups using metadata generated by the LLM pipeline. Datasets containing fewer than two samples passing alignment quality control in either group were excluded. Surrogate variable analysis (sva v3.58.0) was used to estimate latent technical confounders, which were incorporated as covariates in the design matrix to account for unwanted variation and generate batch-adjusted expression results (Leek 2014). Differential expression analysis was performed using linear models with empirical Bayes moderation implemented in limma-voom (v3.66.0), producing log2 fold changes and Benjamini-Hochberg-adjusted p-values (Law et al. 2014). As an alternative approach for gene prioritisation, characteristic direction signatures were calculated from limma-voom normalised expression values (Clark et al. 2014).

### Pathway enrichment

Over-representation analysis of Gene Ontology (GO) terms was performed using the clusterProfiler R package (v4.16.0) (Yu et al. 2012). Analyses were conducted separately for the Biological Process (BP) and Molecular Function (MF) ontologies. Input perturbed genes were provided as Entrez Gene identifiers and GO terms containing between 10 and 500 annotated genes were tested. Statistical significance was assessed using the hypergeometric test against a background of all known human genes, with P values adjusted for multiple testing using the Benjamini-Hochberg false discovery rate (FDR) method.

### Signature comparisons

Query signatures were compared with every signature in the perturbation atlas using one of two similarity measures, chosen by the form of the query. Where the query was a full ranked profile (internal benchmark), similarity was calculated using the cosine similarity between the query and reference score vectors, taken over all genes measured in both. Reference signatures were scored on the characteristic direction statistic or the π-value (−log10 adjusted p × log2 fold change) (Samart et al. 2021) for limma-voom. For up- and down-regulated gene sets queries (external benchmark), similarity was computed by gene set enrichment analysis using fgsea (v1.36.2) (Subramanian et al. 2005) as the difference between the enrichment scores of the query’s up- and down-regulated sets in the ranked reference signature. Similarity scores were sign-inverted where the perturbation directions of query and reference were discordant e.g. overexpression vs knockdown. Performance was assessed as the rank of the best-scoring signature of the correct perturbed gene, after excluding all signatures from the query’s own GEO accession. For each query, 50 genes were randomly sampled in place of the true perturbed gene for comparison.

### Data and software availability

The manually curated training and validation dataset, held-out test dataset, and full production annotation dataset are available on Hugging Face at https://huggingface.co/datasets/jsoul/geo-perturbation-grouping-train. Finetuned model weights for Qwen3-4B-Instruct and Qwen3-30B-A3B-Instruct are available on Hugging Face at https://huggingface.co/jsoul/geo-perturbation-grouping-qwen3-30b-a3b and https://huggingface.co/jsoul/geo-perturbation-grouping-qwen3-4b.

The Nextflow pipeline for automated GEO perturbation annotation is available at GitHub (https://github.com/soulj/LLM-GeoSig). The R package for perturbation signature search and matching is available at GitHub (https://github.com/soulj/perturbMatch). Code to reproduce the figures and analyses presented in this manuscript is available at GitHub (https://github.com/soulj/perturbMatchPaper).

## Results

### Identification of candidates for screening

A keyword search of NCBI-GEO identified 13,061 RNA-Seq and 5,947 microarray experiments with potential gene perturbations. The number of RNA-Seq datasets deposited in GEO has grown substantially in recent years, highlighting the growing challenge of manually identifying and curating suitable perturbation experiments (Figure 1). To automate this process, we developed a Nextflow-based LLM pipeline that identifies case-control sample groupings representing single-gene perturbations and extracts the associated metadata required for downstream differential expression analysis, including the perturbed gene and perturbation type (e.g. knockdown) (Figure 2).

**Figure 1.**
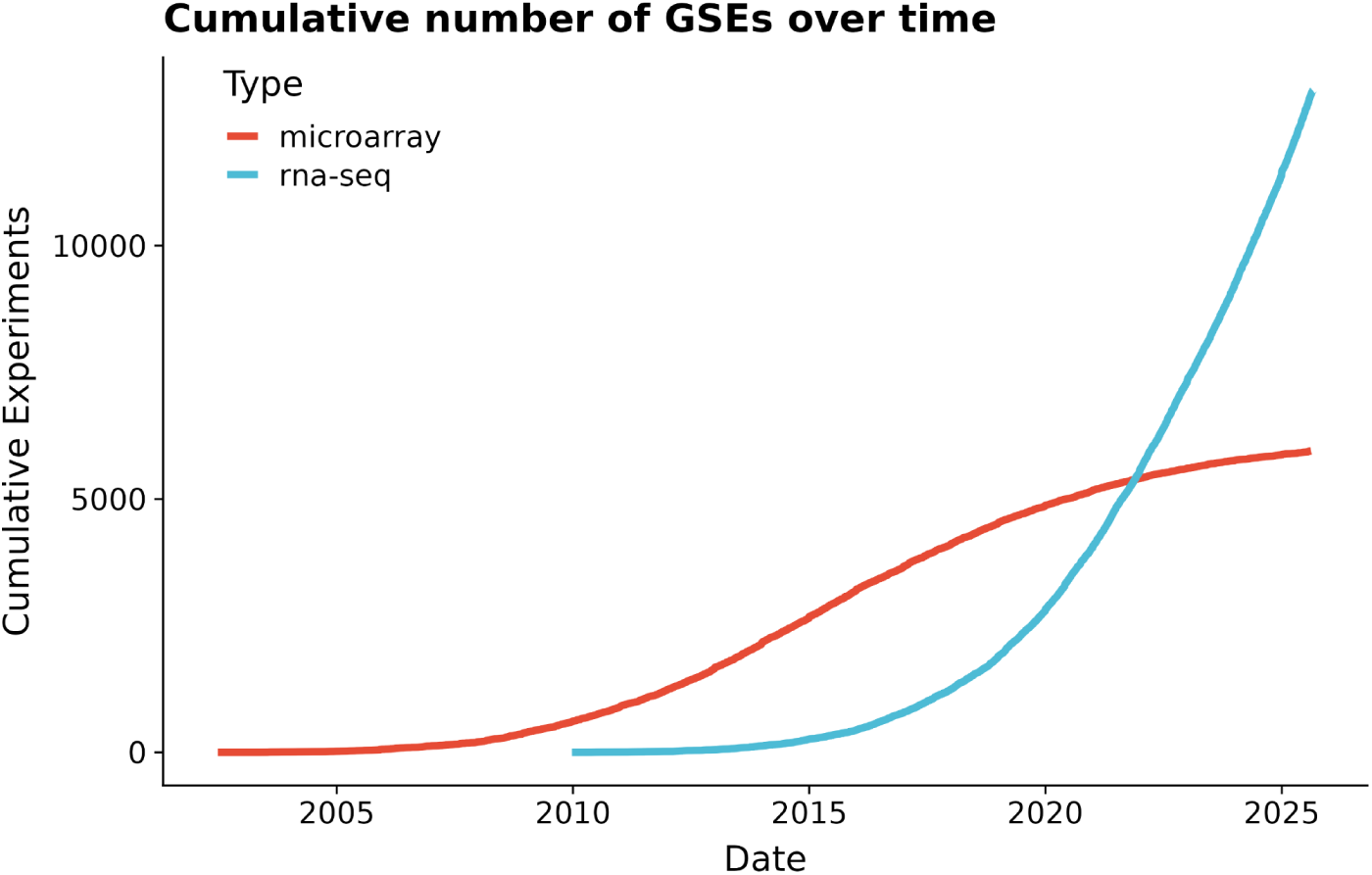
Cumulative number of candidate gene perturbation experiments identified in NCBI-GEO. NCBI-GEO was queried using gene perturbation-related keywords to identify candidate experiments for downstream curation. The cumulative numbers of RNA-Seq and microarray experiments are shown over time.

**Figure 2.**
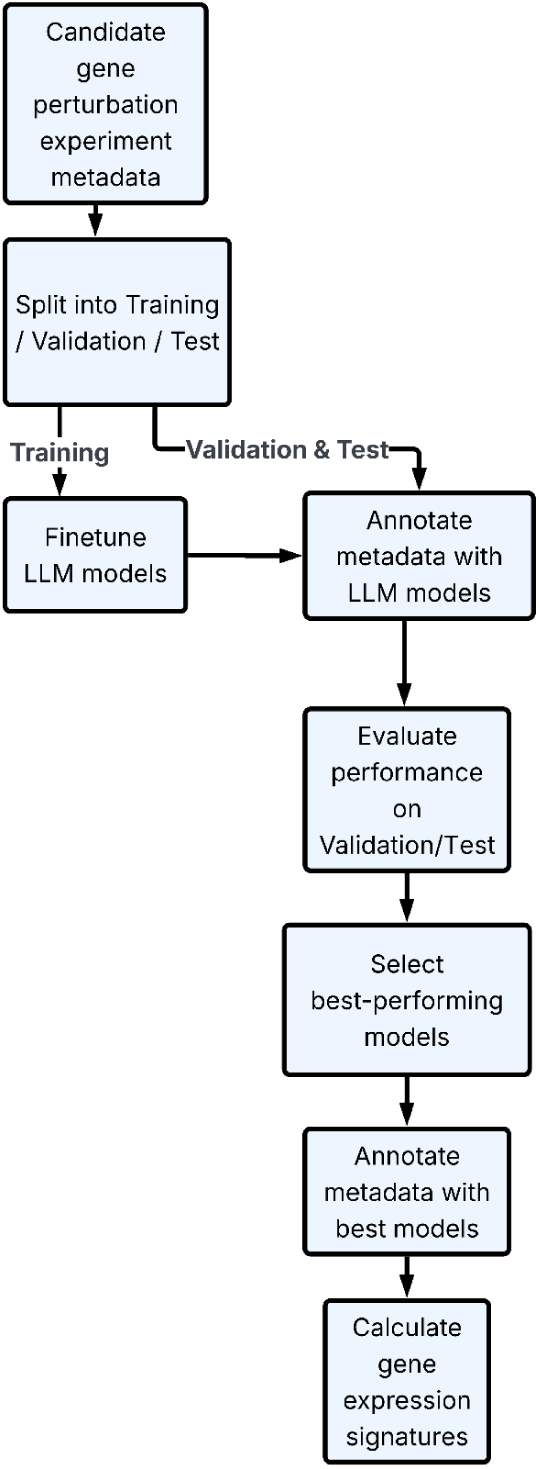
Overview of the LLM driven workflow for automated annotation of GEO gene perturbation experiments. Candidate human gene perturbation studies were identified from GEO and a subset manually curated to define known perturbation-control groups and associated metadata. Curated experiments were partitioned into training, validation, and independent (post-2023) test datasets then used to finetune and evaluate open-weight LLMs for metadata annotation. The best-performing models were applied to annotate the full dataset, enabling generation of gene perturbation expression signatures from the identified perturbation and control sample groups.

### Evaluating performance of LLMs for case-control group prediction

To benchmark local, open-weight LLMs and support model finetuning, we manually curated 3,000 GEO accessions released before 2023, comprising 1,000 RNA-Seq experiments, 1,000 microarray experiments, and 1,000 experiments containing no identifiable perturbation groups. These were randomly split into training (80%) and validation (20%) datasets. This dataset was subsequently used for model finetuning and evaluation.

Across baseline models, validation precision ranged from 0.29 (Gemma-3-4B) to 0.87 (Qwen3-8B) (Table 1, Figure 3A). Within model families, performance generally improved with increasing parameter count e.g. precision increased from 0.29 for Gemma-3-4B to 0.68 for Gemma-3-27B. Reasoning models, including Qwen3-8B and Magistral-Small, consistently outperformed instruction-tuned models of comparable size.

**Figure 3.**
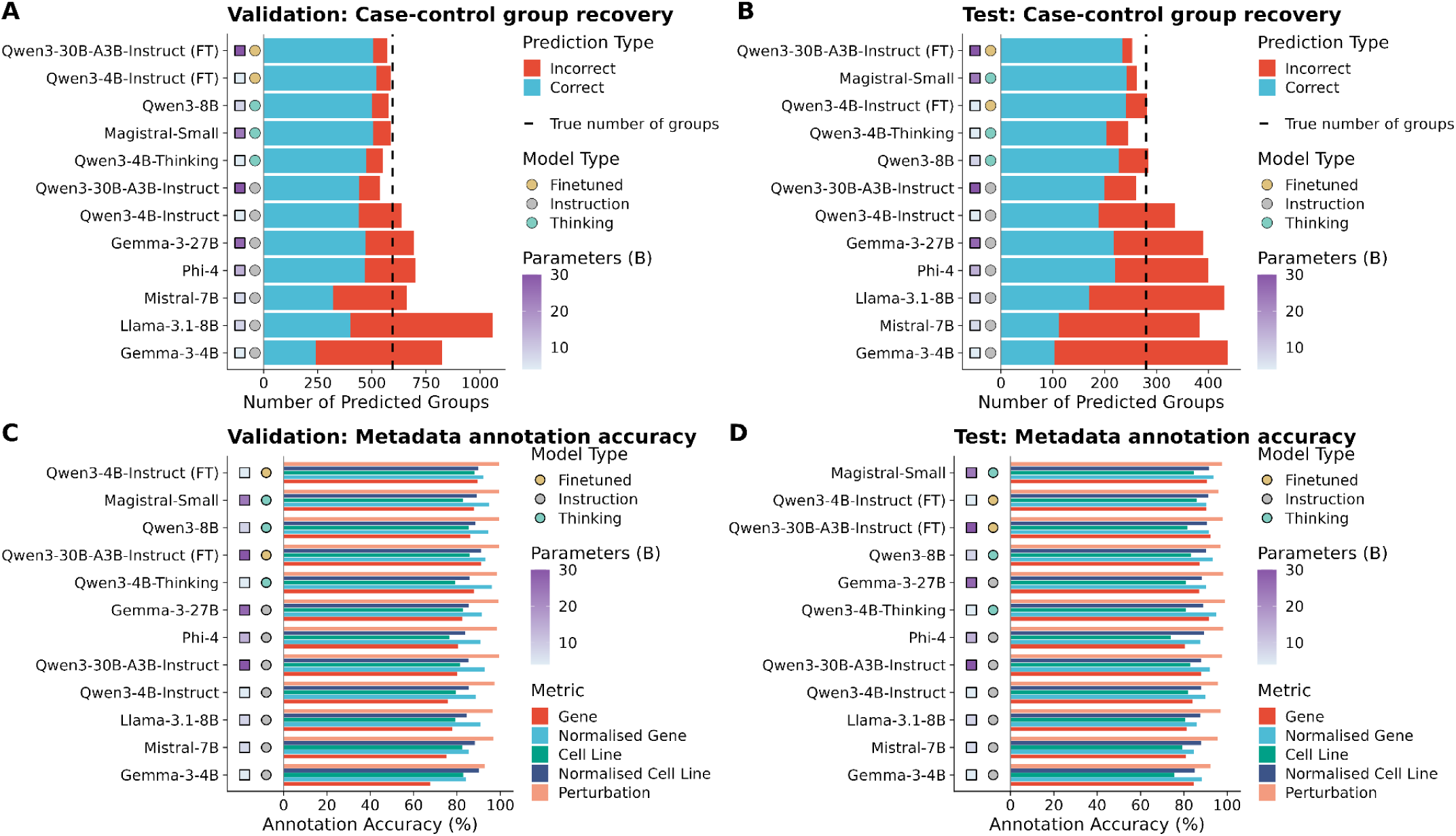
Performance of large language models in reconstructing case-control groupings and annotating metadata. Case-control group recovery on the validation set (A) and the held-out post-2023 test set (B). Stacked bars show the total number of groups predicted by each model; blue indicates groups matching a ground-truth group, red indicates incorrect predictions. The dashed line denotes the true number of groups. Higher-performing models recover more ground-truth groups while making fewer incorrect predictions. (C, D) Metadata annotation accuracy across correctly predicted groups on the validation set (C) and test set (D), for the extracted perturbed gene symbol (Gene), the gene symbol after HGNC normalisation (Normalised Gene), the extracted cell line (Cell Line), the cell line after EFO normalisation (Normalised Cell Line), and the perturbation type (Perturbation). Square fill colour indicates model size (parameters, billions) and circles indicate model type. Models are ordered by performance within each panel. Finetuned models (FT).

**Table 1.** LLM performance on case-control group identification. Precision, recall and F1 score on validation and held-out test sets

| Model | Validation |  |  | Test (post-2023) |  |  |
| --- | --- | --- | --- | --- | --- | --- |
|  | Precision | Recall | F1 | Precision | Recall | F1 |
| Qwen3-30B-A3B-Instruct (FT) | **0.888** | 0.850 | 0.869 | **0.925** | 0.836 | 0.878 |
| Magistral-Small | 0.862 | 0.850 | 0.856 | 0.924 | **0.864** | **0.893** |
| Qwen3-4B-Instruct (FT) | 0.886 | **0.876** | **0.881** | 0.855 | 0.861 | 0.858 |
| Qwen3-4B-Thinking | 0.862 | 0.797 | 0.828 | 0.833 | 0.729 | 0.777 |
| Qwen3-8B | 0.865 | 0.840 | 0.853 | 0.799 | 0.811 | 0.805 |
| Qwen3-30B-A3B-Instruct | 0.823 | 0.741 | 0.780 | 0.762 | 0.711 | 0.736 |
| Qwen3-4B-Instruct | 0.690 | 0.738 | 0.713 | 0.560 | 0.671 | 0.610 |
| Gemma-3-27B | 0.676 | 0.790 | 0.729 | 0.556 | 0.775 | 0.648 |
| Phi-4 | 0.665 | 0.785 | 0.720 | 0.550 | 0.786 | 0.647 |
| Llama-3.1-8B | 0.378 | 0.672 | 0.484 | 0.394 | 0.607 | 0.478 |
| Mistral-7B | 0.487 | 0.541 | 0.513 | 0.290 | 0.396 | 0.335 |
| Gemma-3-4B | 0.292 | 0.405 | 0.339 | 0.236 | 0.368 | 0.287 |

Finetuning the highest-performing non-reasoning models, Qwen3-4B-Instruct and Qwen3-30B-A3B-Instruct, on the curated training dataset increased precision by 0.196 and 0.065, respectively, producing the highest-precision models overall. These results suggest that task-specific finetuning substantially improves case-control group prediction. While several base instruction-tuned models achieved moderate to high recall (0.74-0.79), the finetuned Qwen3 models and Magistral-Small combined recall ≥0.85 with high precision, resulting in the highest overall F1 scores.

To evaluate generalisation to newly deposited data, models were assessed on an independent test set of 300 randomly selected candidate gene perturbation experiments released after 2023. This temporal hold-out was designed to reduce potential bias in the training data and to account for changes in GEO metadata and sequencing technologies over time. The test set comprised 171 RNA-Seq, 10 microarray, and 119 “no group” experiments.

Performance on the held-out test set was similar to that observed during validation, with finetuned and reasoning models again outperforming instruction-tuned models (Figure 3B). The finetuned Qwen3-30B-A3B-Instruct model achieved the highest precision (0.925), while Magistral-Small achieved the highest recall (0.864) and F1 score (0.893). The best-performing models maintained high recall on the independent test set (0.836-0.864), suggesting the pipeline generalises well.

Error analysis was performed on the test dataset for the Magistral-Small and Qwen3-30B-A3B-Instruct (FT) models (Supplementary Table 1). Incorrect application of the exclusion criteria caused models to discard an entire experiment because an excluded term appeared somewhere in its description, despite an allowable subset of samples being present. Precision errors arose when models returned a group mistaken for a perturbation, such as reporter-sorted populations or conditioned media, or paired a valid perturbation with the wrong control samples. A small number of errors reflected erroneous source metadata, e.g. genotype fields with potential spreadsheet autofill errors (GSE222598).

### Metadata identification and normalisation

Following identification of case and control groups, accurate extraction and normalisation of the perturbed gene and mode of perturbation is essential for downstream use of these data, for example retrieval of experiments perturbing a gene of interest. Assessment of exact HGNC symbol matches in the validation data identified cases where the gene name was correctly extracted from the text but did not correspond to a current official symbol. To incorporate up-to-date gene nomenclature into the pipeline, extracted gene symbols were fuzzy-matched against current HGNC symbols and synonyms, with the top five candidates presented to the LLM alongside experiment metadata to disambiguate between closely related synonyms.

This approach improved gene symbol normalisation accuracy across all models, increasing accuracy from 0.68-0.91 using raw extracted symbols to 0.84-0.96 after normalisation (Table 2, Figure 3C, 3D). The largest improvements were observed for lower-performing instruction-tuned models, including Qwen3-4B-Instruct (0.76 to 0.89), Qwen3-30B-A3B-Instruct (0.80 to 0.93) and Llama-3.1-8B (0.78 to 0.91). Following normalisation, gene symbol extraction accuracy was consistently high across models, exceeding 0.94 for the best models. Qwen3-4B-Thinking achieved the highest normalised gene accuracy (0.960).

**Table 2.** Metadata extraction accuracy within correctly identified groups. Proportion of correctly identified groups with accurate metadata on the validation and held-out test **sets**

|  | Validation |  |  | Test (post-2023) |  |  |
| --- | --- | --- | --- | --- | --- | --- |
|  | Gene | Cell line | Perturbation | Gene | Cell line | Perturbation |
| Qwen3-30B-A3B-Instruct (FT) | 0.931 | **0.910** | **0.996** | 0.915 | 0.905 | 0.979 |
| Qwen3-8B | 0.946 | 0.886 | 0.996 | 0.934 | 0.901 | 0.969 |
| Qwen3-30B-A3B-Instruct | 0.927 | 0.853 | 0.995 | 0.920 | 0.881 | 0.975 |
| Qwen3-4B-Instruct (FT) | 0.921 | 0.899 | 0.994 | 0.905 | 0.913 | 0.959 |
| Magistral-Small | 0.949 | 0.892 | 0.994 | 0.938 | **0.917** | 0.975 |
| Gemma-3-27B | 0.915 | 0.855 | 0.991 | 0.903 | 0.881 | 0.982 |
| Phi-4 | 0.908 | 0.838 | 0.985 | 0.877 | 0.894 | 0.982 |
| Qwen3-4B-Thinking | **0.960** | 0.858 | 0.983 | **0.951** | 0.890 | **0.990** |
| Qwen3-4B-Instruct | 0.886 | 0.855 | 0.973 | 0.899 | 0.880 | 0.957 |
| Mistral-7B | 0.854 | 0.883 | 0.969 | 0.847 | 0.879 | 0.955 |
| Llama-3.1-8B | 0.907 | 0.844 | 0.965 | 0.859 | 0.876 | 0.971 |
| Gemma-3-4B | 0.842 | 0.901 | 0.929 | 0.883 | 0.851 | 0.922 |

Cell line annotations were similarly normalised to the EFO ontology to yield standardised cell line identifiers (Methods). Normalised cell line accuracy was high and consistent across models, ranging from 0.84 to 0.91 on the validation set and 0.85 to 0.92 on the held-out test set (Table 2), with the two models used for final annotation, Magistral-Small and finetuned Qwen3-30B-A3B-Instruct, among the best performing (0.89 and 0.91 on validation; 0.92 and 0.91 on test, respectively).

Perturbation type extraction was similarly accurate, ranging from 0.92 to 0.996 and exceeding 0.98 for most models, with Qwen3-30B-A3B-Instruct (FT) and Qwen3-8B performing best. Metadata fields were therefore annotated more reliably than case and control groups were identified, indicating that metadata extraction is the less challenging of the two tasks. Together, these results show that combining LLM extraction with gene normalisation substantially improves robustness to outdated gene symbols and synonym usage, while perturbation type can be identified reliably across a wide range of model architectures.

### Automated atlas of gene perturbations

To generate an atlas from these curated metadata, case-control groups identified from 4,826 manually curated and automatically annotated datasets were processed from NCBI pre-processed count data where available using a limma-based and characteristic direction based differential expression pipeline. Only RNA-Seq experiments were carried forward to the atlas, as uniformly pre-processed intensity matrices are not available for the microarray series. This yielded 6,802 expression signatures from 4,453 experiments, covering 2,907 uniquely perturbed genes. Knockdown signatures (52%) were considerably more common than overexpression (21%) or knockout (26%) signatures (Figure 4B).

**Figure 4.**
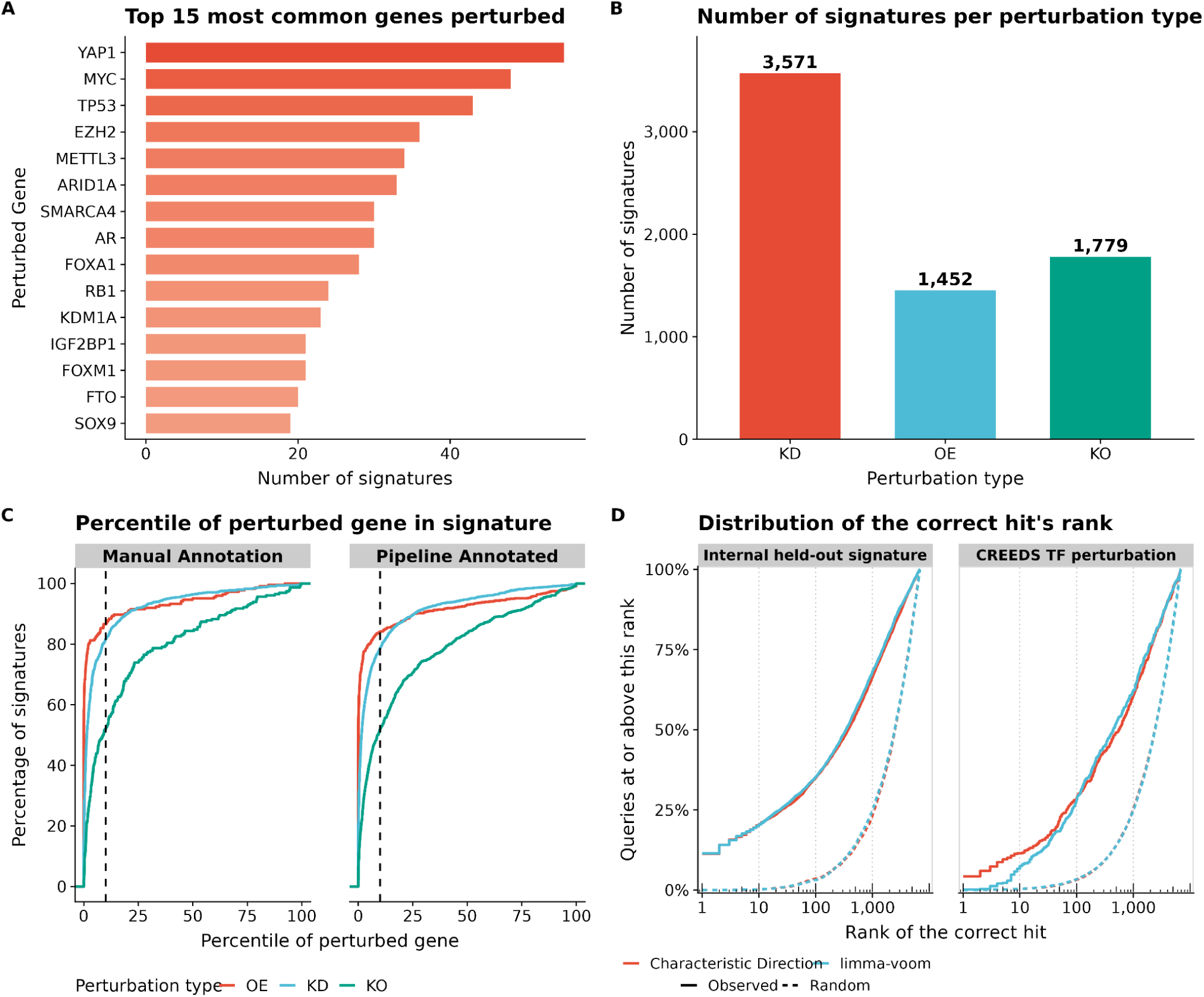
Characteristics and validation of the gene perturbation atlas. (A) Top 15 most frequently perturbed genes in the atlas. (B) Number of expression signatures derived from knockdown (KD), overexpression (OE), and knockout (KO) experiments. (C) Percentile rank of the perturbed gene within its corresponding differential expression signature for manually curated and pipeline-annotated datasets; dashed lines indicate the top 10% threshold. (D) Recovery of known perturbation targets across Characteristic Direction and limma-voom signature databases, for internal held-out signatures (left) and external CREEDS transcription factor perturbations (right). Higher recovery rates indicate improved retrieval of signatures generated by perturbation of the same gene.

Of the perturbed genes, 77 had been targeted in more than 10 experiments, indicating that a subset of genes including YAP1, MYC and TP53 have been studied repeatedly (Figure 4A). To characterise which genes researchers have chosen to perturb we performed functional enrichment analysis. Perturbed genes were enriched for Gene Ontology Biological Process terms including gland development, cell fate commitment, regulation of mRNA metabolic process and stem cell differentiation, and for the Molecular Function term DNA-binding transcription factor binding, consistent with an enrichment of transcription factors among the studied targets (Supplementary Table 2).

As a quality control measure, we examined the rank of the perturbed gene within each signature, considering the expected direction of change in its expression. The perturbed gene was differentially expressed within the top 10% of genes in 78.3% of manually curated signatures, compared with 72.4% of automatically curated signatures, consistent with the performance reported in the benchmarking above. Overexpression perturbations showed stronger differential expression of the target gene overall than knockdown, potentially reflecting incomplete knockdown or off-target effects. Only 52% of knockout signatures, in both the curated and LLM-annotated datasets, had the perturbed gene within the top 10%, consistent with frameshifts that abolish protein function while leaving transcript levels largely intact (Figure 4C).

### R package for signature search

To facilitate exploration and use of this resource, we implemented several signature-matching algorithms in an accompanying R package, perturbMatch. The package supports both atlas exploration and user-driven queries, allowing researchers to submit their own differential expression signatures to identify the most similar perturbation experiments in the atlas e.g. to identify candidate genes whose perturbation mimics or reverses an observed transcriptional response.

Performance was benchmarked using a leave-one-out recovery of known gene perturbations. Each signature was queried against the remainder of the atlas by cosine similarity, excluding all signatures from the same experiment (Figure 4D). The perturbed gene was recovered within the top 1,000 of 6,802 signatures for 66.4% of queries using characteristic direction signatures and 68.1% using limma-derived signatures. External validation using 586 CREEDS microarray transcription factor perturbation gene sets from 191 transcription factors was performed using GSEA scoring (Figure 4D). The perturbed transcription factor was recovered within the top 1,000 for 60.8% of queries using characteristic direction signatures and 62.3% using limma-derived signatures. Performance was comparable between the limma and characteristic direction databases in both benchmarks, suggesting the choice of differential expression approach has limited impact on downstream signature matching.

## Discussion

We have developed and evaluated an LLM-driven pipeline for automated annotation of gene perturbation experiments in NCBI-GEO and applied it at scale to thousands of RNA-Seq datasets. The resulting perturbation atlas enables analyses that would otherwise require substantial manual effort, including identification of existing perturbation signatures that match newly generated gene expression responses. Inconsistent and incomplete metadata annotation across public omics repositories remains a major barrier to data reuse and reproducibility (Rung and Brazma 2013). By enabling scalable, automated extraction of experimental context and biological annotations, this approach improves the accessibility and interoperability of public transcriptomic resources, supporting the broader goals of Findable, Accessible, Interoperable, and Reusable (FAIR) data sharing (Wilkinson et al. 2016).

Reasoning models, including Qwen3-4B-Thinking and Magistral-Small, demonstrated stronger performance on the case-control identification task than comparably sized non-reasoning models. In addition to improved accuracy, these models provide explicit reasoning traces that can aid interpretation of errors and support iterative refinement of prompts and curation strategies. Although larger, closed frontier models may achieve higher raw accuracy, we restricted our evaluation to open-weight models of up to 30B parameters, as these better suit a scalable, updatable resource, offering full reproducibility, independence from third-party API availability or pricing, and archivable weights for future re-annotation. Finetuning produced the largest improvements for smaller models, suggesting that task-specific training is particularly beneficial when base model performance is limited (Chen et al. 2025). Finetuning is comparatively time and resource-intensive relative to few-shot prompting, requiring curated training data and GPU time that must be weighed against the accuracy gains it provides. As LLM capabilities continue to develop rapidly, the curated benchmark dataset generated here provides a resource for systematic evaluation and comparison of future models.

Despite strong performance, LLM-generated annotations should be interpreted with appropriate caution, particularly when applied in high-throughput analyses. For applications where annotation accuracy is critical, users should manually verify case-control assignments and perturbed gene identities as errors in gene identification and grouping of samples remain. Our quality control approach provides an indication of annotation confidence, although highly related genes or genes sharing similar biological functions may also produce strong quality control signals and so require additional validation. Therefore, the pipeline is best viewed as a scalable annotation framework that reduces manual effort, but still necessitates expert review for high-confidence biological interpretation.

Direct benchmarking against existing resources such as CREEDS and the Gene Perturbation Atlas is complicated by differences in curation scope. Those resources encompass a broader range of perturbations, including drug treatments, multi-gene perturbations, and mixed experimental designs, whereas our approach focuses specifically on single-gene perturbation experiments with explicit case-control comparisons. This narrower scope produces annotations that are directly suitable for computing single gene perturbation differential expression signatures. Our resource provides perturbed gene identity and perturbation type as structured metadata fields, which are not systematically captured in broader automated approaches such as RummaGEO.

As with all resources derived from public datasets, the atlas reflects the biases present in GEO and in the broader study of gene function. A relatively small number of well-characterised genes are represented by many perturbation experiments, whereas most human protein-coding genes have been perturbed infrequently or not at all. Large-scale initiatives aiming to systematically characterise the function of all human genes will provide additional perturbational data that can be readily incorporated into the R package and downstream analyses (Adli et al. 2025).

The current pipeline could be improved in several ways, including optimisation of prompting and normalisation strategies for non-coding RNA perturbations, such as lncRNA and circRNA studies (Ramilowski et al. 2020). The framework could similarly be extended beyond GEO to other repositories such as BioStudies or to non-human species. This, however, would likely require additional training data to capture repository-specific metadata conventions and prompt or normalisation strategies that generalise reliably beyond human gene symbols. A strength of the pipeline is its flexibility as the initial GEO search strategy and LLM prompts can be adapted to prioritise specific disease contexts, drug perturbations, or alternative assay types, such as ATAC-seq (Fang et al. 2026). The Nextflow implementation enables updating the database as new GEO datasets become available, while the gene symbol normalisation database can be refreshed as HGNC annotations change without requiring model retraining (Di Tommaso et al. 2017). As additional training data become available and LLM capabilities improve, further prompt refinement and targeted finetuning can extend the scope and accuracy of the resource.

A potential future direction is the integration of linked publications associated with GEO experiments into the annotation pipeline, enabling extraction of methodological and experimental details from manuscripts to confirm or resolve ambiguous metadata. An agent-based framework could selectively invoke literature retrieval and analysis when metadata are incomplete or conflicting, while avoiding additional computational cost for well-annotated experiments. More broadly, embedding LLM-based quality control within the GEO submission workflow could help identify annotation inconsistencies at the point of deposition, improving metadata completeness and promoting more standardised reporting practices.

Gene expression signature databases have emerged as complementary approaches to pathway enrichment methods, with the advantage of directly comparing observed transcriptional responses to perturbations rather than relying on predefined pathway annotations (Lamb et al. 2006; Duan et al. 2020; Schubert et al. 2018). Our R package implements multiple signature similarity approaches, enabling scalable annotation and comparison of perturbational signatures from public transcriptomic data. This resource expands the ability to reuse existing experiments for functional discovery and hypothesis generation.

## Author contributions

**Jamie Soul:** Conceptualisation; Methodology; Software; Investigation; Data curation; Validation; Visualisation; Funding acquisition; Writing - original draft; Writing - review and editing. **David A. Young:** Data curation; Validation; Supervision; Funding acquisition; Writing - review and editing.

